# Microfluidic Nano-Plasmonic Imaging Platform for Purification- and Label-Free Single Small Extracellular Vesicle Counting

**DOI:** 10.1101/2025.03.17.643807

**Authors:** Omid Mohsen Daraei, Avinash Kumar Singh, Saswat Mohapatra, Mohammad Sadman Mallick, Abhay Kotnala, Wei-Chuan Shih

## Abstract

Tumor-derived circulating small extracellular vesicles (sEVs) is a promising non-invasive biomarker for disease diagnosis. However, their quantitative detection remains challenging due to their small size and the complexity of blood plasma. Typically, sample preparation like purification is required. This study presents a purification-free approach using a microfluidic chip integrated with PANORAMA (Plasmonic nano-aperture label-free imaging) for label-free single sEV counting in plasma. CD63, CD9, and CD81 antibodies, specific biomarkers for most sEVs, are functionalized on AGNIS (arrayed gold nanodisks on invisible substrate) for selective capture. The automated microfluidic platform minimizes manual errors and allows precise programming of flow rates, directions, and media for optimization. Only 20 µL of plasma is required, and the analysis is completed within 60 minutes. This platform shows great potential as a sensitive and efficient tool for detecting circulating sEVs without purification or labeling.

## Introduction

Small extracellular vesicles (sEVs) such as exosomes play a role in intercellular communication locally and non-locally via blood circulation ^1–4^. sEVs are not only present in the blood but in virtually all other biological fluids. sEVs are of endosomal origin and their size range is between 30 to 150 nm with an average diameter of around 100 nm ^5^.

The surface proteins and cargo contained within sEVs indicate their cellular origin, making them a valuable subject of study for understanding biological processes ^6,7^. Precise identification and measurement of sEVs are essential for exploring their role in normal physiological processes and disease-associated mechanisms ^8,9^.

Conventional sEV assays such as Western blot, ELISA (enzyme-linked immunosorbent assay), and bead-based flow cytometry have limitations when it comes to assessing the heterogeneity of individual sEV ^10–16^. Additionally, these methods require relatively large volumes of biofluid samples for each assay and are not well-suited for smaller sample sizes ^17–20^.

Nanoparticle tracking analysis (NTA) can count individual sEV and has a size limit on the smaller size end. More importantly, these techniques typically require prior isolation and/or purification which are time-consuming and yield low recovery rates and insufficient purity, thus making their translation toward clinical applications challenging ^21–23^.

Reátegui et al. have introduced an analytical microfluidic platform known as the EVHB-Chip. This platform incorporates engineered nanointerfaces that enable rapid isolation of tumor-specific extracellular vesicles (EVs) in 3 hours, achieving a remarkable 94% specificity for tumor-derived EVs and a limit of detection as low as 100 EVs/ml using fluorescence labels ^24^.

In contrast, plasmonic sensing techniques, particularly the AGNIS (arrayed gold nanodisk on the invisible substrate) platform, provide label-free sEV detection and can be easily combined with fluorescence-based imaging to profile both surface and cargo biomarkers within sEVs ^25,26^. AGNIS consists of high-density gold nanodisk arrays fabricated on a glass substrate with a strategically designed undercut structure achieved by etching away a portion of the glass substrate beneath the gold nanodisks. This innovative structure amplifies localized surface plasmon resonance (LSPR) by enhancing the confinement of electric fields, thereby providing improved sensitivity for detecting single sEVs even in complex biological environments. This method leverages the LSPR of metallic nanostructures to detect sEVs with high sensitivity, eliminating the need for purification. Plasmonic sensing allows for the analysis of sEVs in their native state ^27–29^. For instance, Lee’s group has developed the nano-plasmonic exosomes (nPLEX) assay, which relies on periodic nanohole arrays to capture and detect exosomes ^8^. Ibrahim et al. demonstrated label-free quantification of sEV on AGNIS by standard UV-VIS spectroscopy and achieved an analytical limit of detection of ∼100 sEV/μL ^25^. Mallick et al. demonstrated single sEV detection, sizing, and localization using dynamic plasmonic nano-aperture label-free imaging (D-PANORAMA) ^26,30^. Ohannesian et al. developed a method for detecting, sizing, and analyzing sEVs. It combines PANORAMA, a label-free imaging technique, with fluorescence imaging using Cy3-labeled molecular beacons and PKH67 dye for precise detection and profiling of sEVs ^26^.

As mentioned earlier, microfluidic devices have several advantages over traditional fluid handling and can result in an integrated device with multiple functions. Microfluidic devices can naturally handle small amounts of liquid samples in an automated fashion without manual operation and improving reproducibility. Utilizing these devices addresses critical limitations of traditional sEV detection methods, such as large sample volume requirements, extensive purification steps, and limited sensitivity ^31^. By employing a push-pull flow incubation strategy within a microfluidic channel, our system achieves higher sEV capture efficiency and improved detection sensitivity ^32^. In this study, the term ‘label-free’ specifically pertains to the detection approach, which does not rely on external fluorescent labels, while utilizing antibodies to selectively capture sEVs from complex biological samples. The microfluidic design enables precise control of flow rates, flow direction, and media, thereby facilitating optimal assay conditions ^33^. To harvest these benefits, we propose an integrated approach to perform PANORAMA-based single sEV characterization using the microfluidic device as illustrated in Fig. 1a. The microfluidic chip, designed to pneumatically automate the entire sEV capture-assay pipeline, facilitates high-throughput processing, ultimately improving the accuracy of exosome detection. Capture and detection of plasma-derived purified exosomes were first demonstrated in the integrated microfluidic-PANORAMA system ^34^. PANORAMA is capable of detecting and quantifying sEVs at a minimum concentration of 1 × 10^4^ particles/mL, with a detection limit as low as 16.7 attoM ^26^. The combination of AGNIS-based LSPR sensing and microfluidic automation enables rapid, label-free detection of sEVs in complex biological samples.

**Figure 1:**
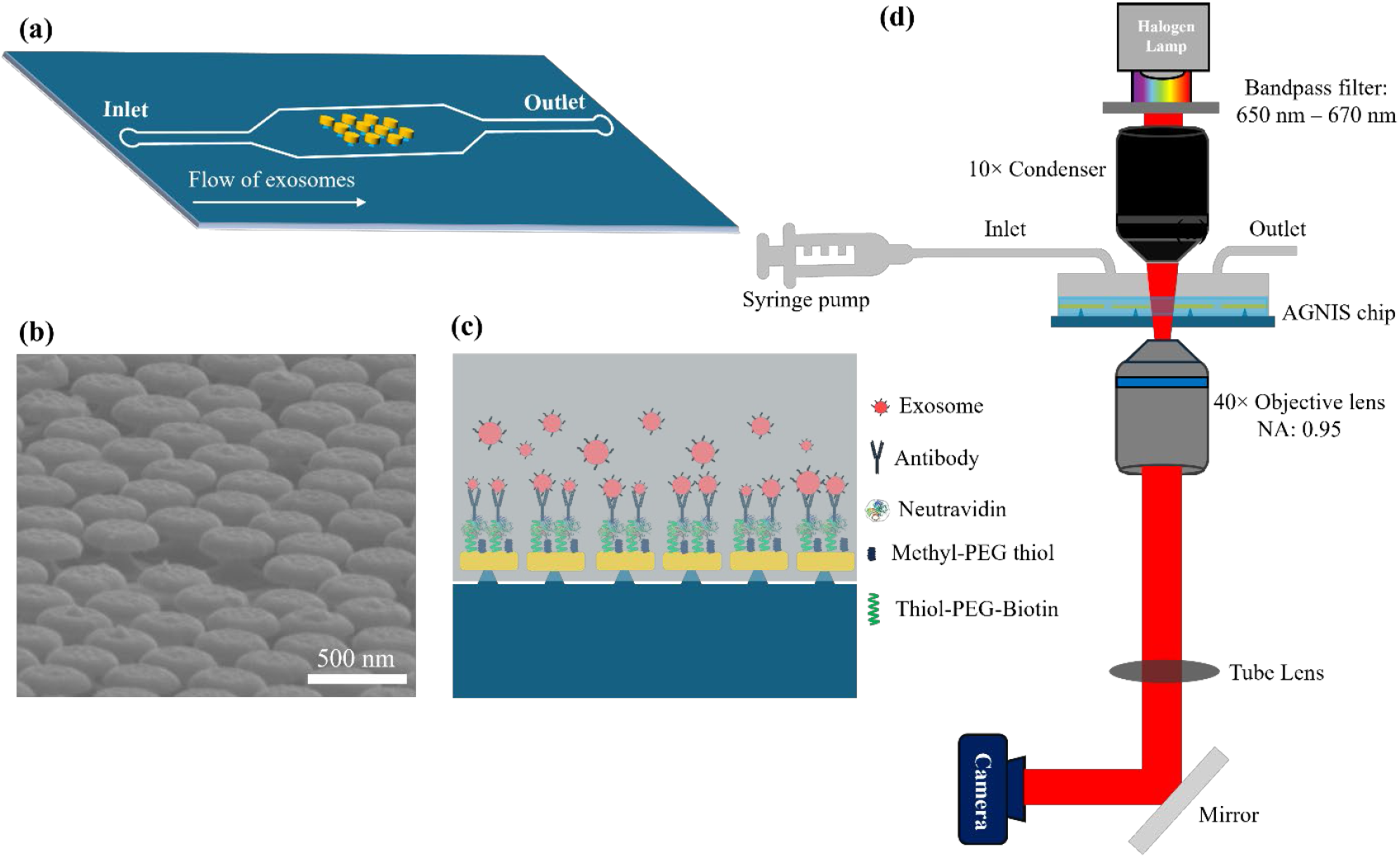
Overview of microfluidic nano-plasmonic imaging platform. (a) Conceptual illustration of the microfluidic device. (b) Scanning electron microscope (SEM) image of the AGNIS surface. (c) Schematic of AGNIS surface functionalization for exosome capture from human plasma. (d) Schematic of the optical setup for the microfluidic-integrated PANORAMA platform.

Advancements in optical detection methods for sEV analysis have further enhanced the capabilities of microfluidic devices. For example, Zhao et al. developed the ExoSearch chip, a microfluidic platform that supports the multiplexed detection of exosomal markers directly from biofluids using LSPR-based optical sensing ^35^. Jhih-Siang Chen et al. also developed a system that combines LSPR with a microfluidic platform, allowing for real-time, label-free detection of sEVs ^36^. These developments demonstrate the utility of optical LSPR sensing within microfluidic devices for sEV analysis.

This study demonstrates the potential of the proposed platform to detect and quantify sEVs and exosomes directly from small plasma volumes without the need for sample purification or labeling, using only 20 µL of plasma from liver cirrhosis patients for the detection and counting of circulating sEVs. The integrated microfluidic platform offers a robust and scalable approach for real-time sEV analysis, supporting various applications in sEV-related research and clinical studies. In this work, the primary focus is on detecting and quantifying sEVs and exosomes, advancing the understanding of their role in biological fluids.

## Results

### LSPR measurement of AGNIS and working principle of PANORAMA

PANORAMA leverages the high refractive index sensitivity of AGNIS’s LSPR to detect nanoparticles. AGNIS shows high refractive index sensitivity due to the radiative coupling of gold nanodisks and an undercut design, exposing regions of high intensity.^25,37–39^ We first measured the LSPR spectrum of the fabricated AGNIS patches and determined their refractive index sensitivity using a custom-built optical setup (See Supplementary Note 1 for details). AGNIS exhibited LSPR peak wavelengths of 659 nm in air and 739 nm in water, yielding a sensitivity of 242.42 nm/RIU, as shown in Supplementary Fig. 1. This high sensitivity has proven effective in detecting nanoparticles via PANORAMA in previous studies.

PANORAMA is a bright-field imaging technique that uses both scattered and unscattered light to detect nanoparticles^30,34^. When the target is far from the AGNIS surface, it behaves like a typical light-scattering object with reduced intensity after transmission through AGNIS. As the target approaches the AGNIS, the local refractive index increases, causing a redshift in the LSPR extinction curve. This redshift decreases extinction and enhances localized light transmission at the operating wavelength range, creating a virtual nanoaperture beneath the target. The increased transmission appears as heightened pixel intensities at the particle’s position on AGNIS. To maximize the signal-to-noise ratio, PANORAMA operates by illuminating with filtered light centered near the LSPR spectrum’s left tail. This enables PANORAMA to detect nanoparticles sub-100 nm nanoparticles, including weakly scattering biological particles, with high sensitivity and precision.

### Purified exosome detection using an integrated microfluidic-PANORAMA platform

To demonstrate the sensing capabilities of our AGNIS-integrated microfluidic device, we first used it to detect purified exosomes extracted from the serum of a liver cirrhosis patient by a standard protocol.^25^ The AGNIS surface was functionalized with exosome-specific antibodies as described in the methods section, to facilitate exosome detection. A 200 µm × 200 µm area of AGNIS, labeled as region 1 in Fig. 6a, was selected as the sensing area within the channel. The channel was initially filled with PBS-1X, and background optical images of AGNIS were acquired. Subsequently, 20 µL of purified exosome solution with a concentration of 7.2×10^5^ exosomes/μL was delivered into the tubing and then flowed into the channel for 1 minute at a flow rate of 5 µL/min. The total channel volume including the inlet and outlet port was ∼5.25 µL. As the channel is filled, exosomes start binding to the AGNIS surface. To enhance binding events, a push-pull incubation strategy was implemented using a pre-programmed syringe pump. In this approach, the exosome solution was alternately moved back and forth within the channel at a flow rate of 1.5 µL/min, with each push and pull cycle lasting for 10 minutes. This method contrasts with the stationary fluid incubation in PDMS wells, where interaction is primarily diffusion driven. It also resolves the issue of depleting a small amount of fluids prior to sufficient exosome-AGNIS interactions with unidirectional flow. The push-pull flow ensures that the entire 20 µL sample solution passes by the AGNIS surface, leading to more binding events and higher particle counts than those with flowless incubation within the channel. Optical images were taken every 20 minutes after the purified exosomes solution was introduced. The push-pull flow was paused temporarily during image acquisition to reduce motion artifacts. The entire process, including sample injection, incubation, and image acquisition, was automated, requiring no manual intervention.

Figure 2a-c shows time-lapsed PANORAMA images of detected exosomes over 60 minutes. After 20 minutes of push-pull incubation, 73 particles were detected, steadily increasing to 537 by the end of 60 minutes (Fig. 2c). This illustrates time-lapsed accumulative capturing of exosomes via antibody-antigen binding. Notably, background noise remained constant at approximately 1% throughout the experiment, demonstrating the microfluidic device’s robustness against external influences compared to open systems. The contrast of exosomes, (Intensity-1) ×100 extracted from the ratioed PANORAMA images was 9.1±1.5 %, as shown in Fig. 2d. The contrast of the exosomes is directly related to their size based on the equation Y=0.153X^2^+7.096X+20.211,^30^ where X % represents the particle contrast. Fig. 2e shows the mean size of the detected exosomes was 99±15 nm obtained from the ratioed PANORAMA image (Fig. 2c).

**Figure 2:**
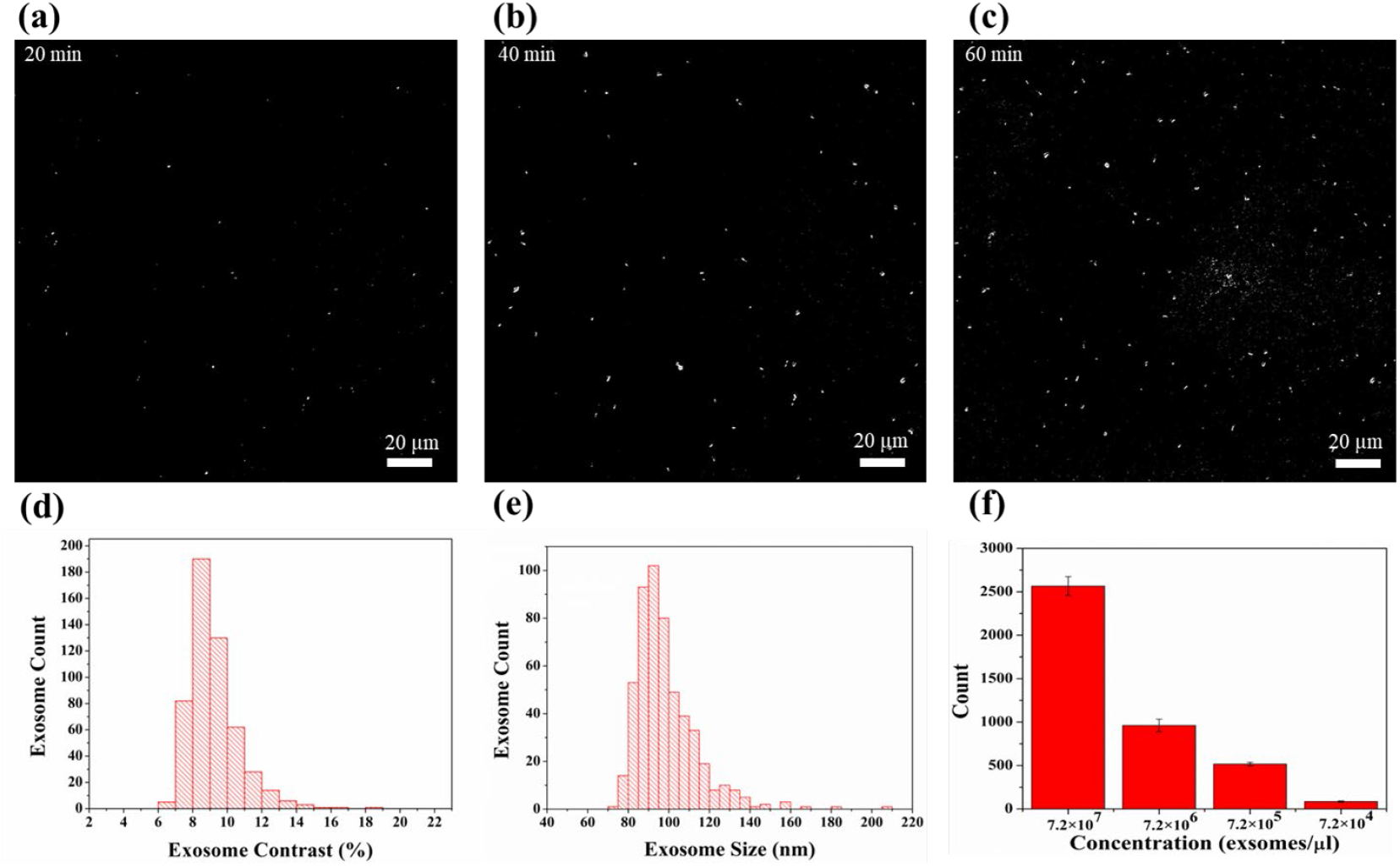
Detection of purified exosome using PANORAMA platform. (a-c) Time-lapse PANORAMA images showing purified exosome detection over 60 minutes, with images captured every 20 minutes. (d) Histogram of the contrast of detected exosomes. (e) Histogram of the size of detected exosomes. (f) Column bar plot representing the average counts and standard deviations for purified exosome samples with four different concentrations, and each concentration was measured in three independent experiments.

Figure 2f displays a column bar plot representing the average exosome counts and standard deviations for purified exosome samples at four different concentrations, ranging from 7.2×10^4^ to 7.2×10^7^ exosomes/µL. The data show a clear trend: as the concentration of exosomes decreases, the mean count of detected exosomes also decreases. Specifically, f or the highest concentration of 7.2×10^7^ exosomes/µL, the mean count is 2565±109, while for the lowest concentration of 7.2×10^4^ exosomes/µL, the mean count drops to 85 ± 7. Intermediate concentrations of 7.2×10^5^ and 7.2×10^6^ exosomes/µL show corresponding mean counts of 516±19 and 962±72, respectively. Each concentration was evaluated through three independent experiments, with exosome counts and contrast values recorded for each trial (See Supplementary Note 2 for details). These results demonstrate the reproducibility of the exosome counts across different concentrations, with consistent measurements observed in three independent experiments for each concentration. The detailed results, including exosome counts and contrast values, are provided in the supplementary information.

### Single sEV detection from patient plasma samples using an integrated microfluidic-PANORAMA platform

Next, we demonstrated the ability of our device to detect and quantify exosomes directly from the plasma of liver cirrhosis patient without any purification. In accordance with MISEV guidelines, we will refer to the detected entities as “sEVs” throughout this section.^2^ AGNIS functionalized with antibodies CD9/CD63/CD81 was employed for sEV detection, with the sensing strategy relying on antigen-specific capture to differentiate sEVs from other plasma components. Initially a volume 20 µL of human plasma was injected into the tubing and then flown into the channel at a flow rate of 5 µL/min for 1 minute to fill the channel with plasma. Just after the flow stops, images were acquired which serve as the background for processing the PANORAMA images. Push-pull incubation, where plasma alternately flowed back and forth in the channel, was employed. Each cycle involved 10 minutes of plasma flow at 1.5 µL/min, with data collection occurring during brief pauses (∼3 seconds) after every 20 minutes. The experiment lasted 60 minutes. Steadily increasing sEVs binding to the AGNIS surface has been observed throughout the incubation, as evidenced by time-lapsed PANORAMA images (Fig. 3a-c). The number of sEV was 246 at 20 minutes after plasma introduction and increased to 943 at 40 min and 1553 at 60 minutes (Fig. 3a-c). The contrast of the detected EVs was extracted from the PANORAMA images, with Fig. 3e displaying the histogram of the contrast. The average contrast of sEVs was 12±3%. The contrast was used to estimate the diameter of detected EVs based on the equation in Section 3.2. The analysis showed that the mean sEV size was 140 nm, with a standard deviation of 33 nm, as shown in Fig. 3f. PANORAMA utilizes optical contrast analysis to achieve single-particle resolution, where intensity changes are directly linked to particle size. This method enables precise differentiation between single sEVs and larger vesicles. The reliability of this approach is supported by size-contrast calibrations and validated through SEM imaging, which provides additional confirmation of individual vesicles captured on the AGNIS surface (Supplementary Note 3 for details). EVs exhibiting less than 18% contrast or a size smaller than 200 nm were categorized as sEVs, while larger particles were classified as large EVs. Following the 60-minute plasma incubation—referred to as Before Wash (BW)—the AGNIS surface was washed by flowing PBS-1X in the channel at a flow rate of 5 µL/min for 5 minutes, to remove non-specifically bound EVs. After Wash (AW) (Fig. 3d), 893 sEVs were retained on the AGNIS surface, exhibiting a mean contrast of 12.3 ± 1.8 % (Fig. 3g). The washing step removed a majority of unbound large sEVs, with the retained sEVs having a mean diameter of 131.4 nm with a standard deviation of 20.69 nm as shown in Fig. 3h. These findings highlight the capability of the microfluidic-integrated PANORAMA system to efficiently and effectively capture, count, and characterize sEVs in human plasma without the need for purification and labeling. It should be noted that regions 1 and 2, shown in Fig. 6a, of AGNIS were used for detecting sEVs. To investigate the impact of AGNIS region selection on sEV capture efficiency within the microfluidic channel, we also performed sensing in region 2 (200 µm × 200 µm). PANORAMA experiments in region 2 revealed a gradual increase in sEV detection over time. At 20 minutes, 185 sEVs were detected, which increased to 624 at 40 minutes and 1132 at 60 minutes (Fig. 3i-k). The contrast of the detected sEVs in region 2 was 11.7 ± 3.4% (Fig. 3m), which was used to estimate the mean diameter of the detected sEVs as 126.6 nm, with a standard deviation of 36.5 nm as shown in Fig. 3n. After washing (Fig. 3l), 668 sEVs were retained on the AGNIS surface with the mean contrast of 11.2 ± 1.8% (Fig. 3o) having the mean diameter of 119.8 nm with the standard deviation of 19.6 nm (Fig. 3p), showing a reduction in particle count compared to region 1, where 893 sEVs were retained. This suggests that exosome capture efficiency is region-dependent within the microfluidic chip, favoring regions near the inlet. In contrast, PANORAMA in PDMS wells showed no significant region-dependent variation in sEV capture efficiency.

**Figure 3:**
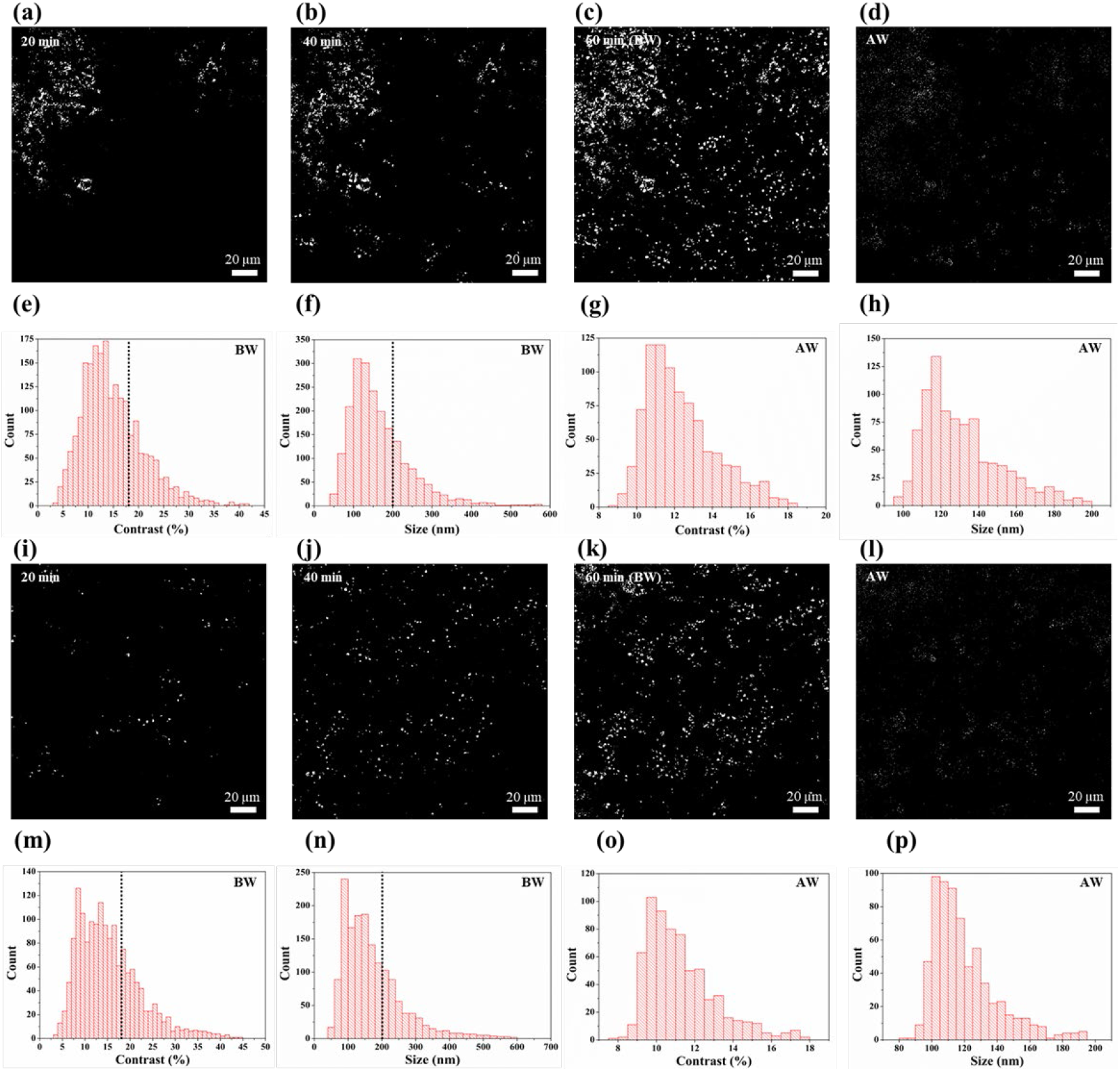
PANORAMA imaging for sEVs detection. Time-lapse PANORAMA images (a-c for region 1, i-k for region 2) illustrate the progressive detection of EVs over 60 minutes (BW), with images captured every 20 minutes. PANORAMA images (d for region 1, l for region 2) show sEVs detection after washing (AW). Histograms of EV contrast (e for region 1, m for region 2) and diameter (f for region 1, n for region 2) during BW are displayed, where the dashed line indicates the cut-off threshold distinguishing sEVs from larger EVs. Similarly, histograms of sEV contrast (g for region 1, o for region 2) and diameter (h for region 1, p for region 2) during AW highlight the size distribution of retained sEVs.

Figure 4 (a) shows the sEVs counts in regions 1 and 2 for BW in three independent experiments, while Figure 4 (b) illustrates the sEVs counts in regions 1 and 2 for AW in these three experiments (See Supplementary Note 4 for details). For BW counts, region 1 exhibits values of 1553, 1451, and 1647 across the three experiments. The mean count for region 1_BW is calculated as 1550 ± 98, indicating stable measurements with moderate variability. In region 2, BW counts of 1132, 1116, and 1183 yield a mean count of 1143 ± 34, showing higher consistency compared to region 1. For AW counts, region 1 displays values of 893, 857, and 905, resulting in a mean count of 885 ± 25, demonstrating low variability across the experiments. In region 2, AW counts of 668, 636, and 689 yield a mean of 664 ± 26, reflecting similar reproducibility as observed for region 1_AW. The relatively low standard deviations across all regions and conditions suggest that the experimental setup provides consistent and reproducible measurements. This consistency highlights the reliability of the method and confirms the robustness of the regions analyzed during the experiments.

**Figure 4:**
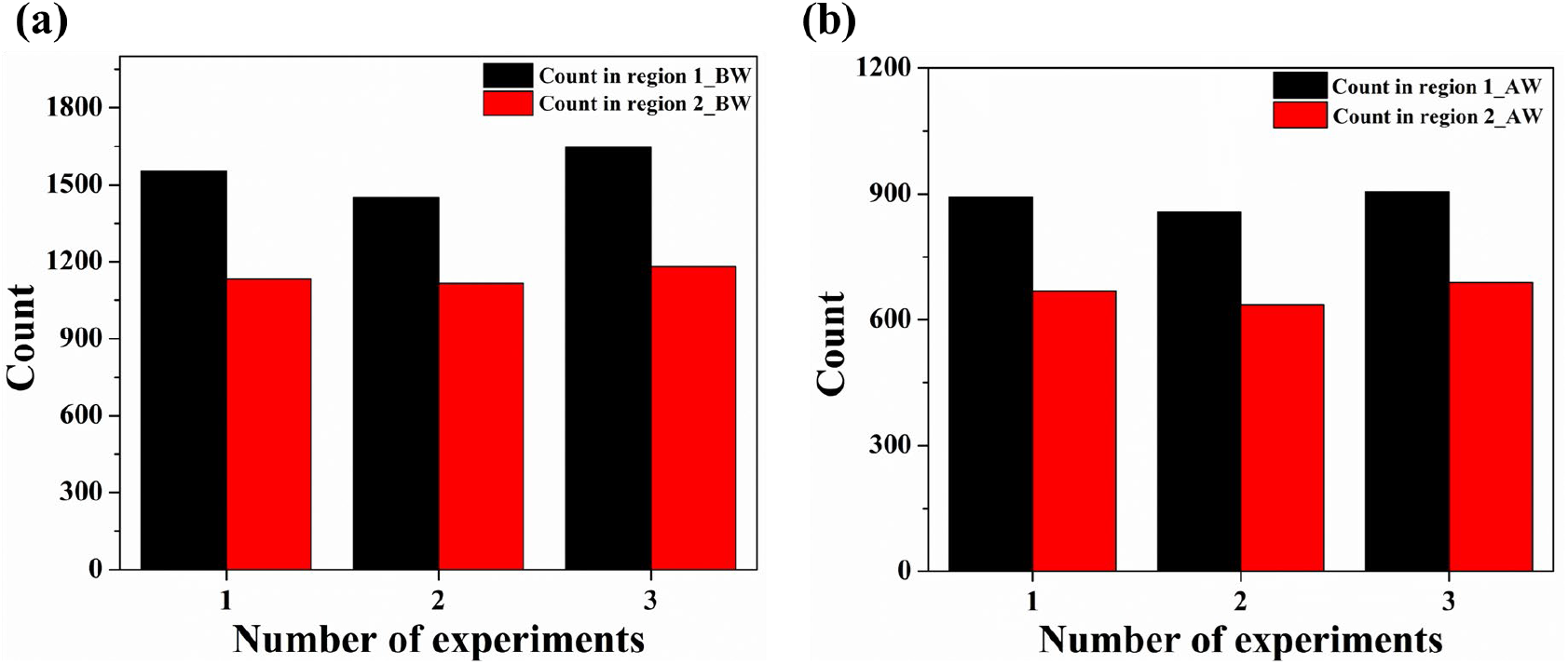
Comparison of BW and AW counts across regions and experiments. (a) Bar plots showing BW counts in region 1 and region 2 across three experiments. (b) Bar plots illustrating AW counts in region 1 and region 2 for the same three experiments.

## Discussion

This study demonstrates the purification- and label-free detection, counting, and size characterization of purified exosomes as well as sEVs in human plasma using the PANORAMA technique integrated with an automated microfluidic system. The use of a push-pull flow incubation strategy within the microfluidic device enhanced the interaction between antibodies and the plasma sample, leading to efficient and effective capture of exosomes/sEVs. With only 20 µL of sample, purified exosomes as well as sEV in human plasma could be captured, detected and counted. The integrated microfluidic-PANORAMA platform, with its minimal sample consumption, purification- and label-free operation, and high detection sensitivity, holds great potential for applications related to exosome/sEV-based fundamental studies and research using animal models.

## Methods

### Reagents

Methyl-PEG-thiol [MT(PEG)4] and bovine serum albumin (BSA) were purchased from ThermoFisher Scientific. Thiol-PEG-Biotin was acquired from Nanocs Inc. Polystyrene beads and neutravidin were purchased from Sigma-Aldrich. Biotinylated antibodies CD9, CD63, and CD81 were obtained from BioLegend. Negative (SU8) and positive (AZ1512) photoresists were procured from Microchem and AZ Electronic Materials, respectively. Krayden Dow Sylgard 184 silicone elastomer kit was purchased from ThermoFischer.

### Optical setup of microfluidic-integrated PANORAMA platform

Optical images of the AGNIS were acquired using a standard inverted optical microscope (IX83, Olympus), as illustrated in Fig. 1d. The AGNIS-integrated microfluidic chip, which was connected to a syringe via tubing and powered by a mechanical syringe pump (Chemyx Inc. Model Fusion 720), was placed on the microscope stage. The AGNIS patch within the microfluidic channel was illuminated by a halogen lamp (U-LH100L3, Olympus) through a 10× condenser lens (IX2-LWUCD, Olympus). To confine the output of the halogen lamp to 650-670 nm, a bandpass filter (FB660-10, Thorlabs) was used. Transmitted light was captured with a 40× objective lens (UPlanSApo 40×/0.95, Olympus) and imaged using a sCMOS camera (Hamamatsu Orca). The exposure time was set to 30 milliseconds, and the imaging area spanned 200 µm × 200 µm.

### Fabrication of AGNIS-integrated microfluidic device

#### Fabrication of AGNIS

AGNIS were fabricated using nanosphere lithography (NSL) as illustrated in Fig. 5. First, glass cover slides (VWR, No. 2, 24×50 mm) were cleaned in acetone, isopropanol, and distilled water followed by drying at 100 °C for 30 minutes. A 2 nm chromium (Cr) adhesion layer and an 80 nm gold (Au) film were then deposited onto the slides using DC sputtering at a rate of 3 Å/s. A monolayer of 460 nm polystyrene beads (PSB) was assembled on the Au surface via the Langmuir– Blodgett technique. The PSB size was reduced to 360 nm using oxygen plasma etching (50 W, 200 V, 30 mTorr), resulting in a closely packed array that served as the mask for the fabrication of nanodisks. Subsequent argon ion milling removed Au between the PSBs. The PSBs were then removed by sonication, yielding a structured array of gold nanodisks (AGN) with a disk diameter of 360 nm and pitch of 460 nm over the entire coverslip.

**Figure 5:**
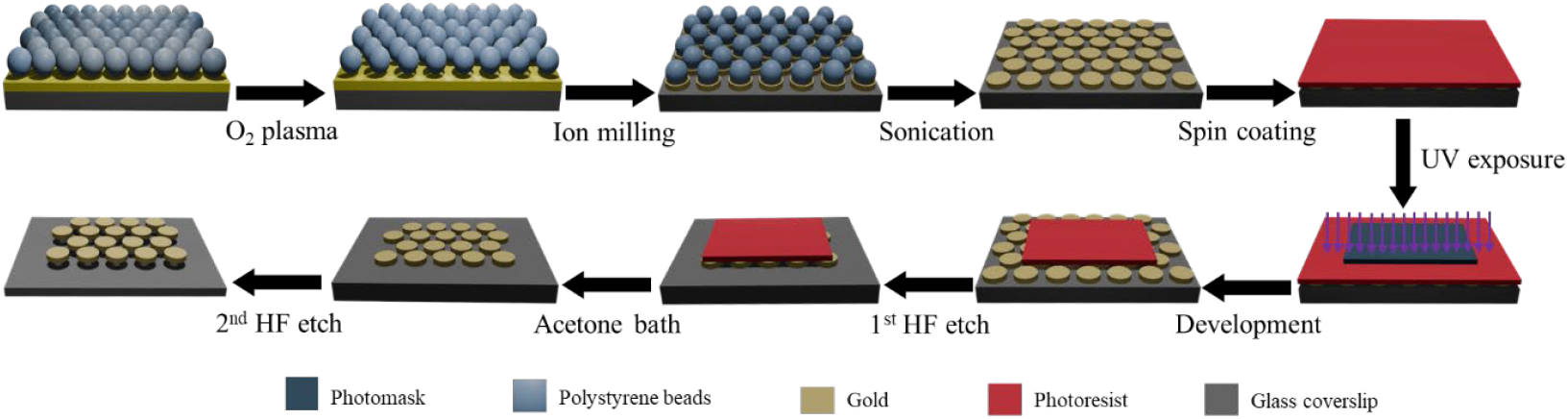
Overview of the fabrication of AGNIS patches. Schematics illustrating the fabrication steps of rectangular AGNIS patches for integration with the microfluidic channel.

To integrate AGNIS into the microfluidic channel, one requires to bond the AGNIS coverslip with the Polydimethylsiloxane (PDMS) block with the channels. However, the presence of AGN across the entire coverslip surface hinders PDMS-glass plasma bonding. To address this, rectangular AGN patches (1000 × 450 µm) were retained on the coverslip, with AGN removed from the remaining areas using photolithography. The AGN surface was uniformly coated with a positive photoresist (AZ1512) and exposed for 3.2 seconds at 80 mJ/cm^2^ with a lamp power of 25 mW/cm^2^, using a mask with rectangular regions. The photoresist was then developed with AZ 300 MIF, exposing the AGN in the unmasked areas. These exposed regions were etched by immersing the chip in buffered hydrofluoric acid (BHF), which removed the underlying glass and the nanodisks. This process enabled effective bonding of the AGNIS chip to the microfluidic channels. After etching, the remaining photoresist was stripped using acetone, leaving behind precisely patterned rectangular AGN regions. The patterned substrate was further immersed in BHF for 80 seconds, to create an undercut beneath the nanodisks, forming AGNIS. This undercut structure enhances electric field confinement and shifts the LSPR peak, thereby increasing the sensitivity for detecting nanoparticles such as sEVs. Fig. 1b shows a representative image of a section of the AGNIS. The undercut results in a blue-shifted LSPR peak that enhances compatibility with visible and near-infrared light sources and detectors, and better diffraction-limited resolution.

#### Fabrication of microfluidic channel and integration with AGNIS

Soft lithography was used to fabricate the microfluidic (MF) channels. A SU-8 mold was created on a silicon wafer using standard photolithography techniques (See Supplementary Note 5 for fabrication details). The design features four identical channels, each with an inlet, outlet, and middle section as illustrated in Fig. 6a. The inlet and outlet sections are identical with a width of 200 µm and length of 4.3 mm. The middle segment is 2 mm in length and 1 mm in width. The middle section is wider than the inlet/outlet to accommodate the AGNIS (1000 × 450 µm) and facilitate manual alignment during bonding. The thickness of the channel was set to 100 µm.

**Figure 6:**
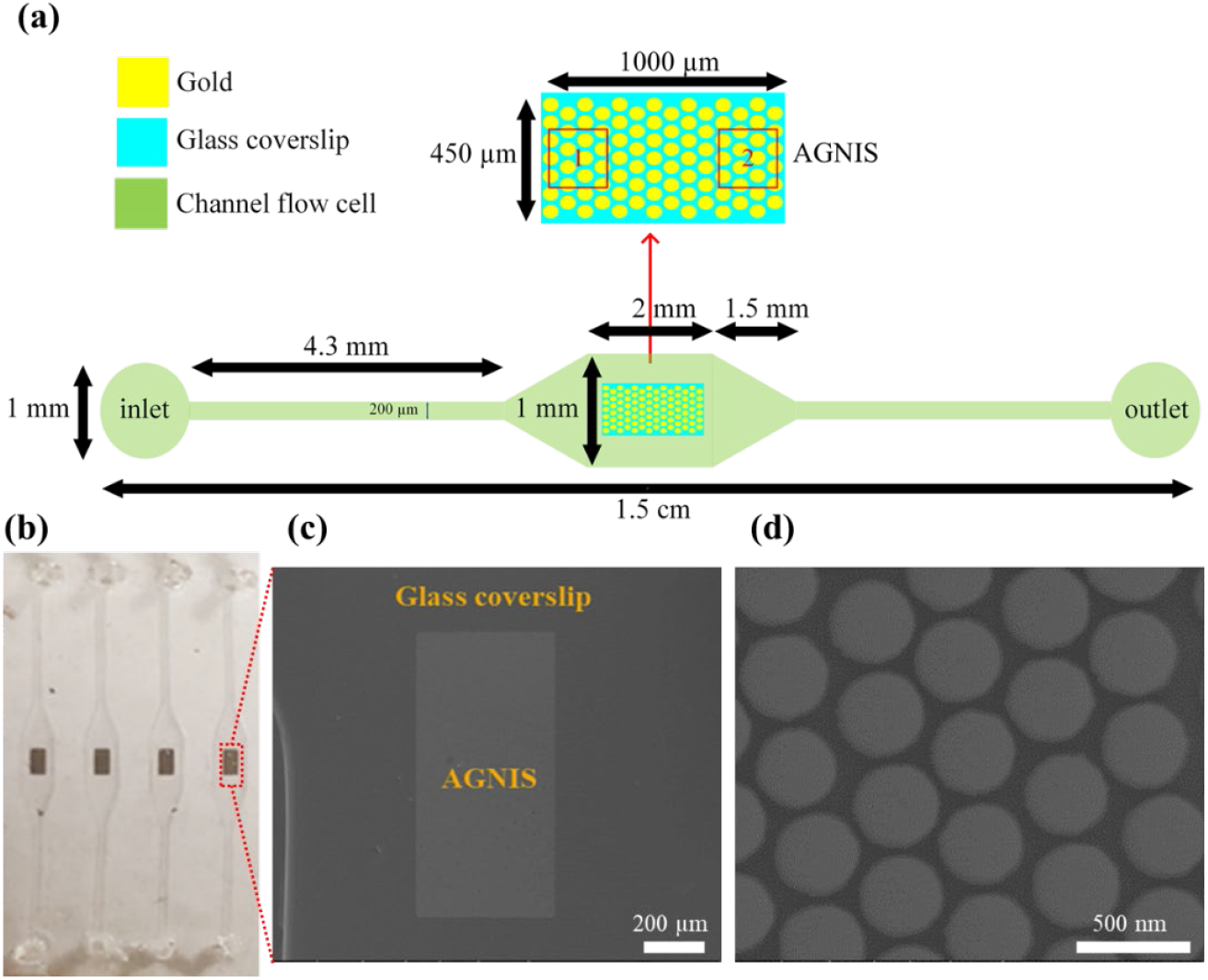
Details of the microfluidic channel. (a) Microfluidic channel design. (b) Optical image of the AGNIS-integrated microfluidic device. (c) SEM image of a single AGNIS patch within the microfluidic channel. (d) SEM image (top view) of densely packed AGNIS from a small area in Figure 6c.

Polydimethylsiloxane (PDMS) was prepared by mixing silicon elastomer and curing agent in a 10:1 ratio, then poured onto the SU-8 mold and cured at 70°C for 2 hours. After curing, the PDMS structure was peeled from the mold, and inlet and outlet channels were punched with a 1 mm biopsy puncher, followed by cleaning with acetone, isopropanol, and deionized water. The microfluidic channels were bonded to the AGNIS chip using oxygen plasma treatment to ensure a strong PDMS-glass bond. This bonding strategy deliberately incorporates small gaps between the AGNIS and the channel walls to ensure strong adhesion and minimize the risk of leakage during operation. Although extending AGNIS coverage might enhance capture efficiency, this approach effectively balances operational functionality with structural stability. Fig. 6b shows the AGNIS-integrated microfluidic device with four channels and rectangular AGNIS patches in the middle section. The SEM image in Fig. 6c and 6d -shows the densely packed nanodisks within the microfluidic channel.

#### Surface functionalization of AGNIS within the microfluidic channel

To functionalize the AGNIS surface within the microfluidic channel, a serial incubation process was employed. First, a 1:3 mixture of long-chain active biotin-PEG-thiol (2mM, MW:1 kDa) and short inactive methyl-PEG-thiol (2mM, MW: 0.2 kDa) was introduced into the channel with a flow rate of 5 µL/min using a syringe pump (Chemyx Inc, Model Fusion 720). The microfluidic chip was then incubated for approximately 16 hours, allowing the thiol groups to bind strongly with the Au nanodisks. After incubation, the channels were washed with phosphate-buffered saline (PBS-1X) with a flow rate of 20 µL/min to remove unbound PEG-thiol molecules. Next, neutravidin with a flow rate of 5 µL/min was introduced as a linker molecule and incubated for 2 hours, followed by thorough washing. Finally, a mixture of exosome-specific antibodies (CD9, CD63, and CD81 with a concentration of 50 µg/mL) diluted with 2.5% BSA is flown through the channel with a flow rate of 5 µL/min and incubated for an additional 2 hours. It should be noted that all the incubations were done at 4 °C. Fig 1c. illustrates the AGNIS surface after functionalization. The AGNIS-integrated microfluidic chip was then used for sEVs detection via antibody-antigen-specific binding.

#### Procedure for plasma extraction

All participants provided consent under institutional review board (IRB)-approved biomarker protocols at MD Anderson Cancer Center. Patients also consented to IRB-approved prospective biomarker or therapeutic protocols based on their disease site. Blood samples were collected via venipuncture before the initiation of treatment (radiation, surgery, or chemotherapy), and the extracted plasma was aliquoted and stored at −80°C until use. The analysis of these samples for sEV research was conducted under protocol 2021-0368, which permits the use of de-identified samples from consented patients enrolled in IRB-approved studies. The approved IRB ID is 2021-0368_CR001, with registration ID IRB 2 IRB00002203.

## Data Availability

All source data supporting this study’s findings are available from the corresponding author upon reasonable request.

## Acknowledgments

This work was supported by the National Institutes of Health (NIH) R01EB-030623 grant. Additionally, the authors thank Dr. Steven H Lin (MD Anderson Cancer Center) for general oncology expertise, Dr. Prasun Jalal (Baylor College of Medicine) and Dr. Manal M. Hassan (MD Anderson Cancer Center) for providing a deidentified patient plasma sample.

## Author Contributions

W.C.S. conceived the idea and directed the study. O.M.D. and W.C.S. designed the experiment and analyzed the data. O.M.D. experimented with the MF device and produced the figures. S.M. fabricated the AGN by NSL, obtained SEM images and O.M.D. fabricated the rectangular AGNIS substrate by photolithography. O.M.D., A.K.S., and S.M. fabricated a PDMS flow cell and attached it to the AGNIS. O.M.D, M.S.M, A.K., and W.C.S. interpreted the data and explained the results. O.M.D., M.S.M., S.M., A.K., and W.C.S. wrote and edited the manuscript

## References

1. Wang, X., Tian, L., Lu, J. & Ng, I. O. L. Exosomes and cancer-Diagnostic and prognostic biomarkers and therapeutic vehicle. Oncogenesis 11, (2022).

2. Théry, C. et al. Minimal information for studies of extracellular vesicles 2018 (MISEV2018): a position statement of the International Society for Extracellular Vesicles and update of the MISEV2014 guidelines. J Extracell Vesicles 7, (2018).

3. Gurung, S., Perocheau, D., Touramanidou, L. & Baruteau, J. The exosome journey: from biogenesis to uptake and intracellular signalling. Cell Communication and Signaling 2021 19:1 19, 1–19 (2021).

4. Liu, W. et al. Insight into extracellular vesicles in vascular diseases: intercellular communication role and clinical application potential. Cell Communication and Signaling 2023 21:1 21, 1–34 (2023).

5. Kalluri, R. & LeBleu, V. S. The biology, function, and biomedical applications of exosomes. Science (1979) 367, (2020).

6. Rajagopal, C. & Harikumar, K. B. The origin and functions of exosomes in cancer. Front Oncol 8, (2018).

7. Kalluri, R. The biology and function of exosomes in cancer. J Clin Invest 126, 1208–1215 (2016).

8. Im, H. et al. Label-free detection and molecular profiling of exosomes with a nano-plasmonic sensor. Nat Biotechnol 32, 490–495 (2014).

9. Ender, F., Zamzow, P., von Bubnoff, N. & Gieseler, F. Detection and Quantification of Extracellular Vesicles via FACS: Membrane Labeling Matters! International Journal of Molecular Sciences 2020, Vol. 21, Page 291 21, 291 (2019).

10. Chen, K. et al. A magneto-activated nanoscale cytometry platform for molecular profiling of small extracellular vesicles. Nature Communications 2023 14:1 14, 1–15 (2023).

11. Jia, Y. et al. Small extracellular vesicles isolation and separation: Current techniques, pending questions and clinical applications. Theranostics 12, 6548–6575 (2022).

12. Dias, T. et al. An electro-optical platform for the ultrasensitive detection of small extracellular vesicle sub-types and their protein epitope counts. iScience 27, (2024).

13. Zhang, J. et al. Extracellular Vesicles: Techniques and Biomedical Applications Related to Single Vesicle Analysis. ACS Nano 17, 17668–17698 (2023).

14. Malhotra, S. et al. Novel devices for isolation and detection of bacterial and mammalian extracellular vesicles. Mikrochim Acta 188, (2021).

15. Yin, H., You, S., Li, X., Li, S. & Guo, C. Progress, challenges, and prospects of small extracellular vesicles isolation and characterization. Journal of Holistic Integrative Pharmacy 5, 121–130 (2024).

16. Davidson, S. M. et al. Methods for the identification and characterization of extracellular vesicles in cardiovascular studies: from exosomes to microvesicles. Cardiovasc Res 119, 45–63 (2023).

17. Zhu, L. et al. Label-free quantitative detection of tumor-derived exosomes through surface plasmon resonance imaging. Anal Chem 86, 8857–8864 (2014).

18. Théry, C. et al. Proteomic Analysis of Dendritic Cell-Derived Exosomes: A Secreted Subcellular Compartment Distinct from Apoptotic Vesicles. The Journal of Immunology 166, 7309–7318 (2001).

19. Zhou, H. et al. Collection, storage, preservation, and normalization of human urinary exosomes for biomarker discovery. Kidney Int 69, 1471–1476 (2006).

20. Lamparski, H. G. et al. Production and characterization of clinical grade exosomes derived from dendritic cells. J Immunol Methods 270, 211–226 (2002).

21. Sourkouni, G., Argirusis, C. & Argirusis, N. Recycling of Surface-Functionalized Nanoparticles—A Short Review. Processes 2024, Vol. 12, Page 2354 12, 2354 (2024).

22. Dilsiz, N. A comprehensive review on recent advances in exosome isolation and characterization: Toward clinical applications. Transl Oncol 50, 102121 (2024).

23. Colao, I. L. Development of a Process Step for the High Purity Recovery of Exosome Material from a Regenerative Cell Product. Doctoral thesis, UCL (University College London). (2021).

24. Reátegui, E. et al. Engineered nanointerfaces for microfluidic isolation and molecular profiling of tumor-specific extracellular vesicles. Nature Communications 2018 9:1 9, 1–11 (2018).

25. Misbah, I., Ohannesian, N., Qiao, Y., Lin, S. H. & Shih, W. C. Exploring the Synergy of Radiative Coupling and Substrate Undercut in Arrayed Gold Nanodisks for Economical, Ultra-Sensitive Label-Free Biosensing. IEEE Sens J 21, 23971–23978 (2021).

26. Ohannesian, N. et al. Plasmonic nano-aperture label-free imaging of single small extracellular vesicles for cancer detection. Communications Medicine 2024 4:1 4, 1–10 (2024).

27. Zhang, P. et al. Molecular and functional extracellular vesicle analysis using nanopatterned microchips monitors tumor progression and metastasis. Sci Transl Med 12, (2020).

28. Wang, J., Trau, M. & Wuethrich, A. A Microfluidic SERS Assay to Characterize the Phenotypic Heterogeneity in Cancer-Derived Small Extracellular Vesicles. Methods in Molecular Biology 2679, 241–253 (2023).

29. Das, S., Lyon, C. J. & Hu, T. A Panorama of Extracellular Vesicle Applications: From Biomarker Detection to Therapeutics. ACS Nano 18, 9784–9797 (2024).

30. Mallick, M. S., Misbah, I., Ohannesian, N. & Shih, W. C. Single-Exosome Counting and 3D, Subdiffraction Limit Localization Using Dynamic Plasmonic Nanoaperture Label-Free Imaging. Adv Nanobiomed Res 3, (2023).

31. Andriolo, G. et al. Methodologies for Scalable Production of High-Quality Purified Small Extracellular Vesicles from Conditioned Medium. Methods in Molecular Biology 2668, 69–98 (2023).

32. Pramanik, S. K. & Suzuki, H. Microfluidic device with a push–pull sequential solution-exchange function for affinity sensing. Microfluid Nanofluidics 23, 1–8 (2019).

33. Pattanayak, P. et al. Microfluidic chips: recent advances, critical strategies in design, applications and future perspectives. Microfluid Nanofluidics 25, 1–28 (2021).

34. Ohannesian, N., Misbah, I., Lin, S. H. & Shih, W. C. Plasmonic nano-aperture label-free imaging (PANORAMA). Nature Communications 2020 11:1 11, 1–10 (2020).

35. Zhao, Z., Yang, Y., Zeng, Y. & He, M. A microfluidic ExoSearch chip for multiplexed exosome detection towards blood-based ovarian cancer diagnosis. Lab Chip 16, 489–496 (2016).

36. Chen, J. S., Chen, P. F., Lin, H. T. H. & Huang, N. T. A Localized surface plasmon resonance (LSPR) sensor integrated automated microfluidic system for multiplex inflammatory biomarker detection. Analyst 145, 7654–7661 (2020).

37. Zhao, F., Arnob, M. M. P., Zenasni, O., Li, J. & Shih, W. C. Far-field plasmonic coupling in 2-dimensional polycrystalline plasmonic arrays enables wide tunability with low-cost nanofabrication. Nanoscale Horiz 2, 267–276 (2017).

38. Dmitriev, A. et al. Enhanced nanoplasmonic optical sensors with reduced substrate effect. Nano Lett 8, 3893–3898 (2008).

39. Acimovic, S. S. et al. Superior LSPR substrates based on electromagnetic decoupling for on-a-chip high-throughput label-free biosensing. Light: Science & Applications 2017 6:8 6, e17042–e17042 (2017).

