## Supplementary file for "Microfluidic Nano-Plasmonic Imaging Platform for Purification- and Label-Free Single Small Extracellular Vesicle Counting"

### Supplementary Note 1:

The optical configuration for acquiring LSPR spectra of AGNIS was constructed using an inverted Olympus microscope (IX71). Illumination of the AGNIS sample, positioned on the sample holder, was achieved with a 100W white halogen lamp, focused through a 0.55 NA condenser. The transmitted light was captured by a 40×/0.75 numerical aperture (NA) objective lens (UplanFLN40X) and directed to a spectrometer (Princeton Instruments, Acton 2300) equipped with a thermoelectrically cooled (-70°C) CCD camera (Princeton Instruments, PIXIS 400) via a 4-f system. A schematic of the optical setup is presented in Supplementary Fig. 1a. AGNIS exhibited LSPR peak wavelengths of 659 nm in air and 739 nm in water, corresponding to a sensitivity of 242.42 nm/RIU (Supplementary Fig. 1b).

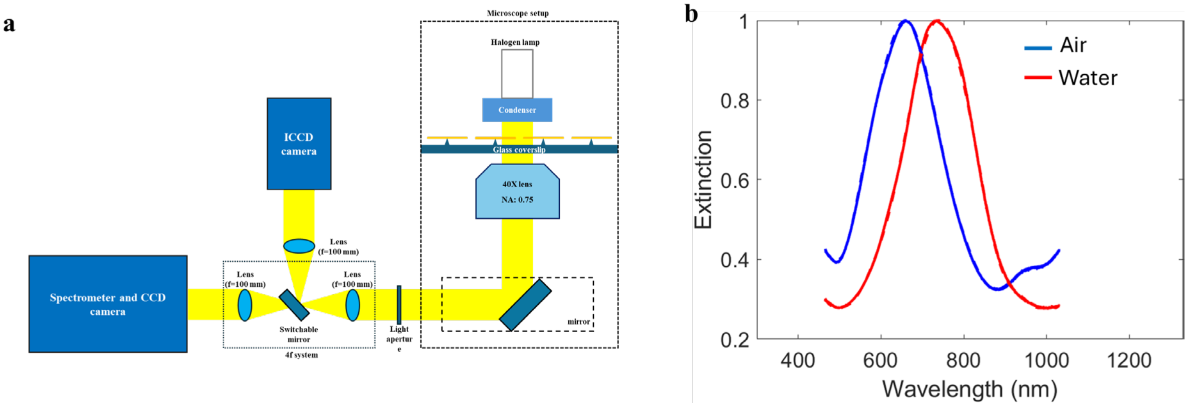


*Supplementary Figure 1: a) Schematic of LSPR imaging system setup, b) LSPR spectra of AGNIS in air and water medium*

### Supplementary Note 2:

Exosome detection was carried out at four different concentrations: $7.2 \times 10^4$ exosomes/µl, $7.2 \times 10^5$ exosomes/µl, $7.2 \times 10^6$ exosomes/µl, and $7.2 \times 10^7$ exosomes/µl. The results of these experiments are detailed below, with references to the corresponding figures for clarity.

At a concentration of $7.2 \times 10^4$ exosomes/µl, the number of detected exosomes in experiments 1, 2, and 3 were 83, 79, and 94, respectively (Supplementary Fig. 2a-c). The corresponding exosome contrasts for these experiments were measured as 10.2 ± 1.7%, 8.3 ± 0.8%, and 9.7 ± 2.2%, respectively (Supplementary Fig. 2d-f).

At a concentration of $7.2 \times 10^5$ exosomes/µl, the number of detected exosomes in experiments 1, 2, and 3 were 537, 515, and 498, respectively (Supplementary Fig. 2g-i). The corresponding exosome contrasts for these experiments were measured as 9.1 ± 1.5%, 8.6 ± 1.7%, and 9.5 ± 1.1%, respectively (Supplementary Fig. 2j-l).

At a concentration of $7.2 \times 10^6$ exosomes/µl, the number of detected exosomes in experiments 1, 2, and 3 were 968, 1031, and 887, respectively (Supplementary Fig. 3a-c). The corresponding exosome contrasts for these experiments were measured as 8.7 ± 1.5%, 9.3 ± 1.9%, and 9.2 ± 1.5%, respectively (Supplementary Fig. 3d-f).

At a concentration of $7.2 \times 10^7$ exosomes/µl, the number of detected exosomes in experiments 1, 2, and 3 were 2563, 2458, and 2676, respectively (Supplementary Fig. 3g-i). The corresponding exosome contrasts for these experiments were measured as 8.7 ± 1.6%, 9.0 ± 2.4%, and 10.8 ± 2.0%, respectively (Supplementary Fig. 3j-l). These results demonstrate the relationship between exosome concentration and detection efficiency, with higher concentrations yielding larger counts of detected exosomes. The measured exosome contrasts remain within a relatively narrow range across different concentrations and experimental repetitions, indicating the reproducibility of the detection process.

**Concentrations: $7.2\times10^4$ and $7.2\times10^5$ $[\frac{\text{exosomes}}{\mu\text{L}}]$**

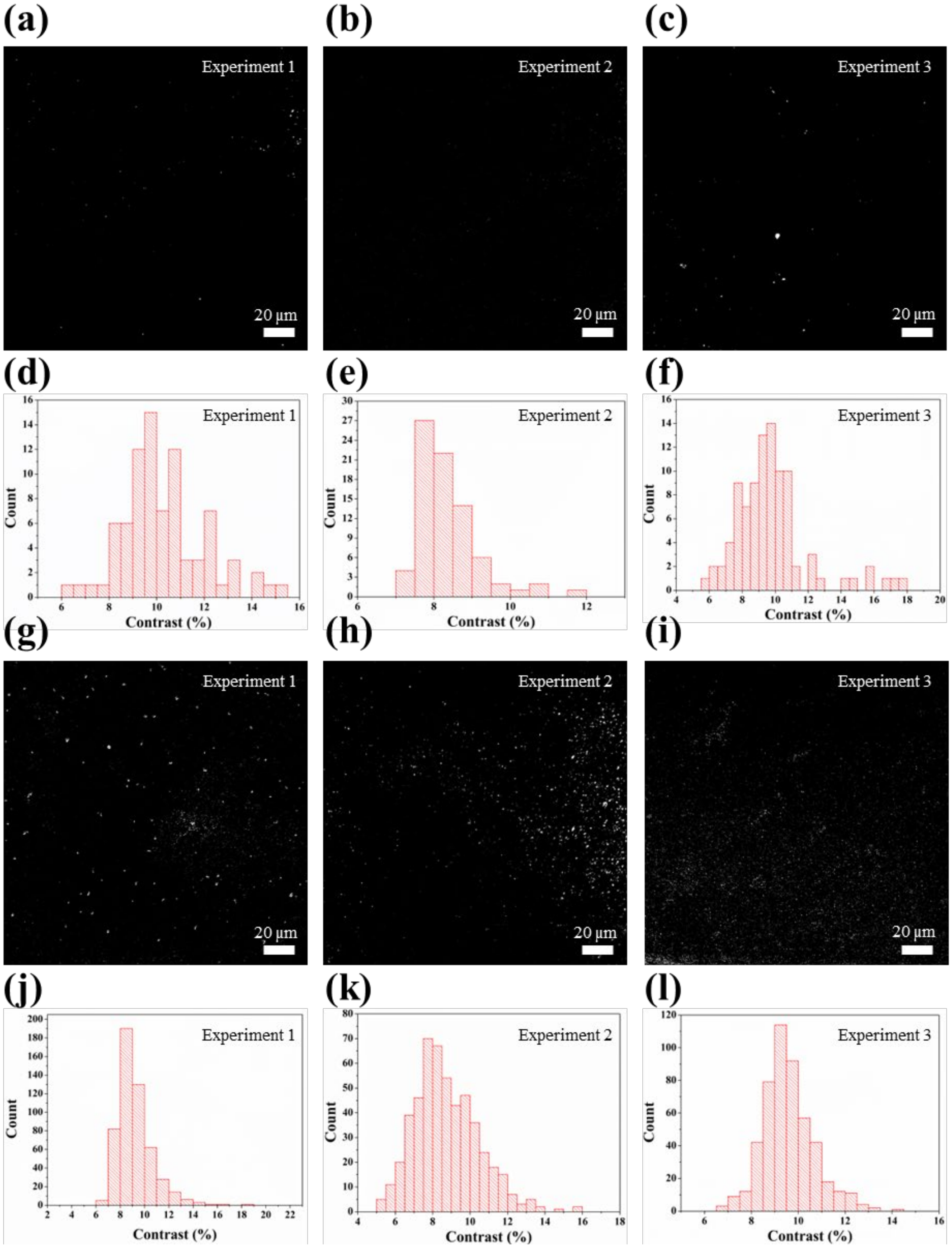


*Supplementary Figure 2: (a-c) PANORAMA images illustrating purified exosome detection from experiments 1 to 3 at concentrations of 7.2 × 10$^4$ exosomes/µl. (d-f) Histograms of the contrast of the detected exosomes from experiments 1 to 3 at the same concentration. (g-i) PANORAMA images illustrating purified exosome detection from experiments 1 to 3 at a concentration of 7.2 × 10$^5$ exosomes/µl. (j-l) Histograms of the contrast of the detected exosomes from experiments 1 to 3 at this higher concentration.*

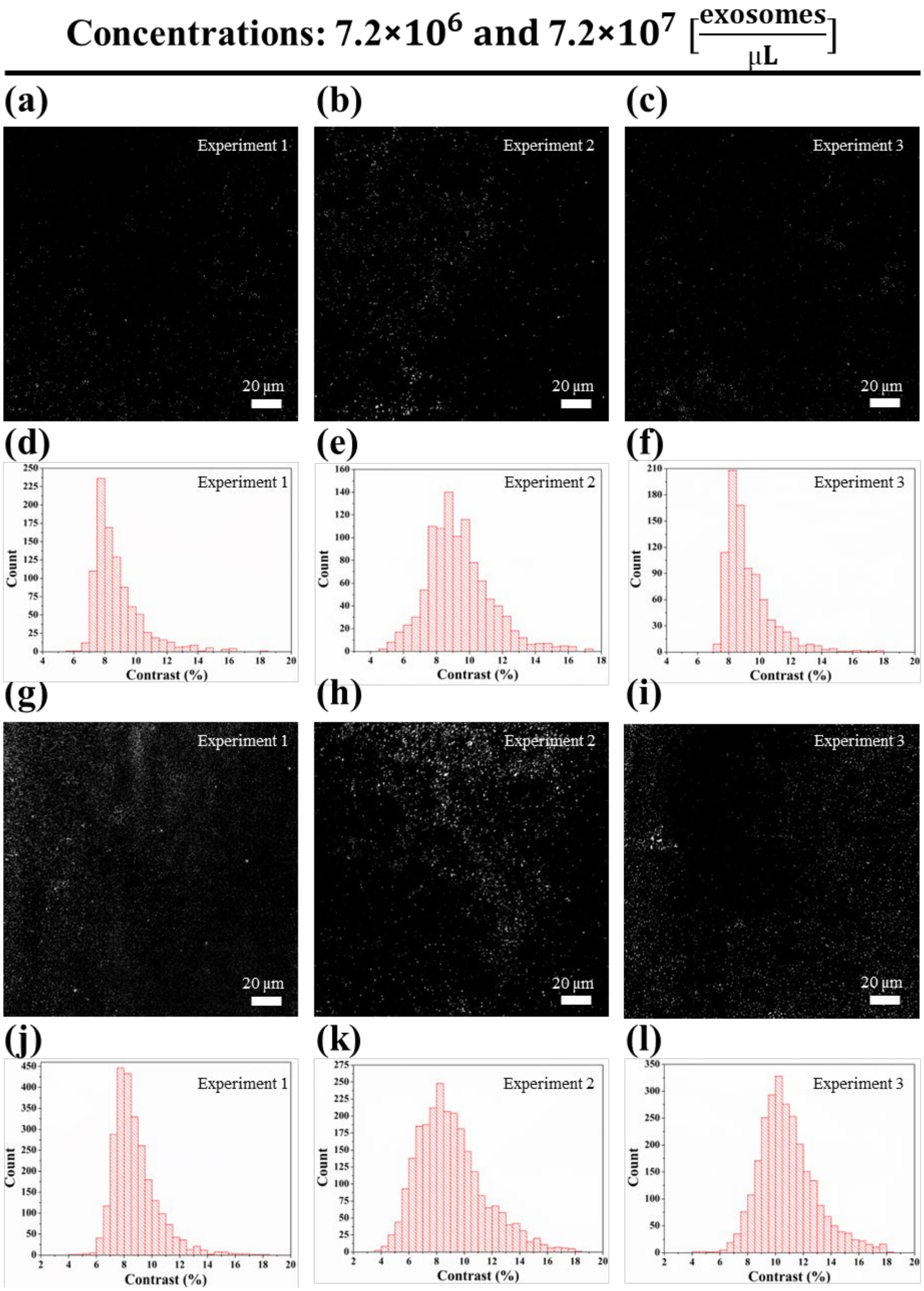


*Supplementary Figure 3: (a-c) PANORAMA images illustrating purified exosome detection from experiments 1 to 3 at concentrations of 7.2 × 10⁶ exosomes/µl. (d-f) Histograms of the contrast of the detected exosomes from experiments 1 to 3 at the same concentration. (g-i) PANORAMA images illustrating purified exosome detection from experiments 1 to 3 at a concentration of 7.2 × 10⁷ exosomes/µl. (j-l) Histograms of the contrast of the detected exosomes from experiments 1 to 3 at this higher concentration.*

### Supplementary Note 3:

The SEM images demonstrate the capability of the AGNIS platform to facilitate and support the attachment of sEVs (Supplementary Fig. 4a,b). The distinct nanoscale features observed in the SEM images confirm the presence of sEVs, providing visual evidence to verify their interaction with the AGNIS surface. The interaction and settlement of sEVs on the AGNIS surface are essential for advancing biological analysis and enhancing the performance of biosensing platforms. The successful settlement observed in this study highlights the effectiveness of the AGNIS substrate in capturing sEVs, demonstrating its potential as a reliable tool for applications in these fields.

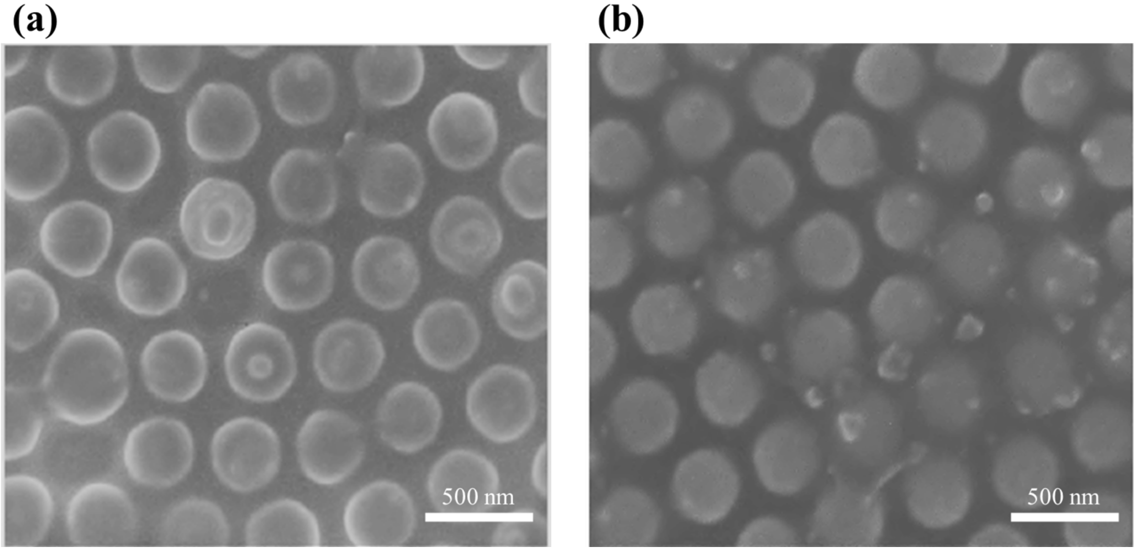


*Supplementary Figure 4: (a,b) SEM images displaying sEVs settlement on AGNIS surface.*

### Supplementary Note 4:

The PANORAMA experiments demonstrated a significant variation in sEV capture efficiency between the two AGNIS regions within the microfluidic channel. To ensure the reproducibility of sEV counts in region 1 and region 2, two additional experiments were conducted in both regions, revealing consistent capture trends and validating the reliability of the system.

In experiment 2, region 1 for BW was analyzed with a threshold of 3.7%, resulting in a count of 1451 sEVs (Supplementary Fig. 5a) and a contrast of 11.4% ± 2% (Supplementary Fig. 5e), equivalent to a mean size distribution of 122.3 ± 22.1 nm (Supplementary Fig. 5f), with a retention rate of 59%. For AW in region 1, a threshold of 3.6% resulted in a count of 857 sEVs (Supplementary Fig. 5b) and a contrast of 10.2% ± 1.9% (Supplementary Fig. 5i), equivalent to a mean size distribution of 109.5 ± 20.7 nm (Supplementary Fig. 5j). In region 2, BW with a threshold of 3.8% recorded a count of 1116 sEVs (Supplementary Fig. 5c) and a contrast of 11.6% ± 1.9% (Supplementary Fig. 5g), equivalent to a mean size distribution of 123.5 ± 21.3 nm (Supplementary Fig. 5h), with a retention rate of 57%. For AW in region 2, a threshold of 3.5% yielded a count of 636 sEVs (Supplementary Fig. 5d) with a contrast of 10.9% ± 1.7%

(Supplementary Fig. 5k), equivalent to a mean size distribution of 116.1 ± 18 nm (Supplementary Fig. 5l).

In experiment 3, region 1 BW had a threshold of 3.8%, resulting in an sEV count of 1647 (Supplementary Fig. 6a) and a contrast of 12.4% ± 2.3% (Supplementary Fig. 6e), equivalent to a mean size distribution of 132.4 ± 25.7 nm (Supplementary Fig. 6f). The retention rate for this region was recorded at 54.9%. For AW in region 1, a threshold of 3.5% produced an sEV count of 905 (Supplementary Fig. 6b) and a contrast of 10.2% ± 1.7% (Supplementary Fig. 6i), equivalent to a mean size distribution of 108.8 ± 18.4 nm (Supplementary Fig. 6j). In region 2, BW with a threshold of 3.9% recorded a count of 1183 sEVs (Supplementary Fig. 6c) and a contrast of 11.6% ± 2% (Supplementary Fig. 6g), equivalent to a mean size distribution of 123.3 ± 21.5 nm (Supplementary Fig. 6h), with a retention rate of 58.24%. Finally, AW in region 2, with a threshold of 3.8%, resulted in a count of 689 sEVs (Supplementary Fig. 6d) and a contrast of 10.3% ± 1.5% (Supplementary Fig. 6k), equivalent to a mean size distribution of 109.9 ± 16.5 nm (Supplementary Fig. 6l). The data highlights the differences in performance between the two methods and the variability across regions. This suggests higher exosome capture efficiency in the AGNIS region near the inlet, likely due to the sample's initial interaction with AGNIS in region 1, where all sEVs in the plasma are available for binding. As the sample flows forward, the availability of sEVs decreases, reducing capture efficiency near region 2. Therefore, region 1 provides optimal conditions for maximizing capture efficiency, making it a preferred choice for sEV detection within the microfluidic channel.

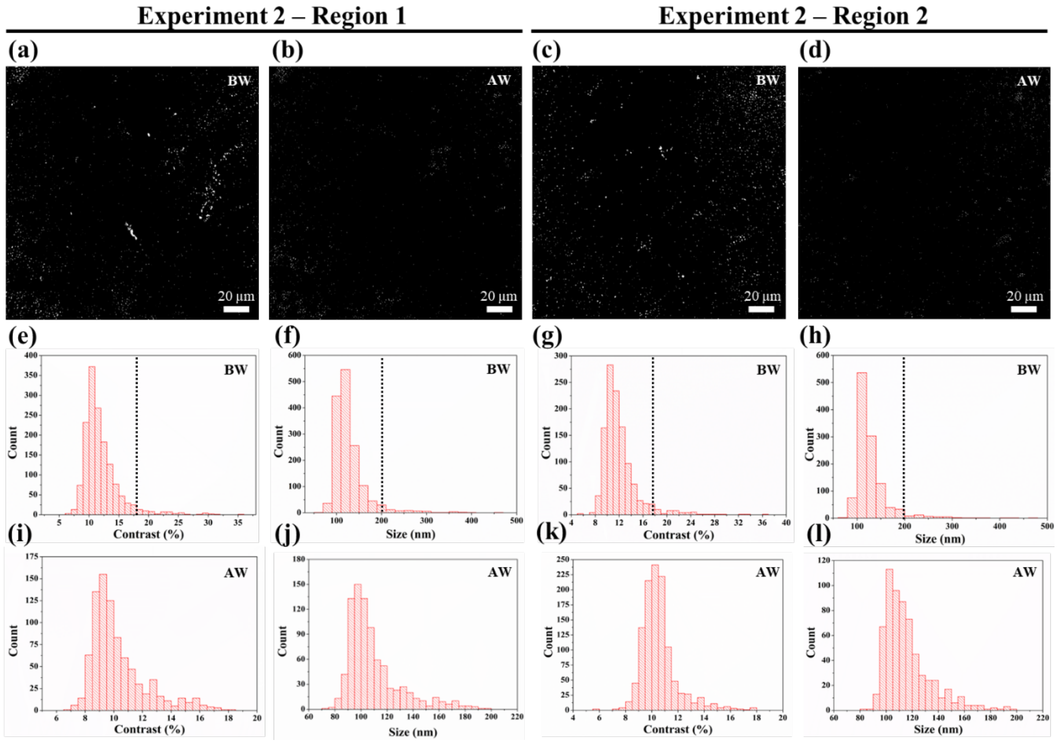


*Supplementary Figure 5: (a, c) PANORAMA images showing EV detection at 60 minutes (BW) in regions 1 and 2 for experiment 2. (b, d) PANORAMA images illustrating sEV detection (AW) in regions 1 and 2 for the same experiment. Histograms of EV contrast during BW are displayed in (e) region 1 and (g) region 2, alongside histograms of EV diameter in (f) region 1 and (h) region 2, with dashed lines indicating the cut-off threshold separating sEVs from larger EVs. Similarly, histograms of sEV contrast during AW are shown in (i) region 1 and*

*(k) region 2, while histograms of sEV diameter are presented in (j) region 1 and (l) region 2, highlighting the size distribution of retained sEVs.*

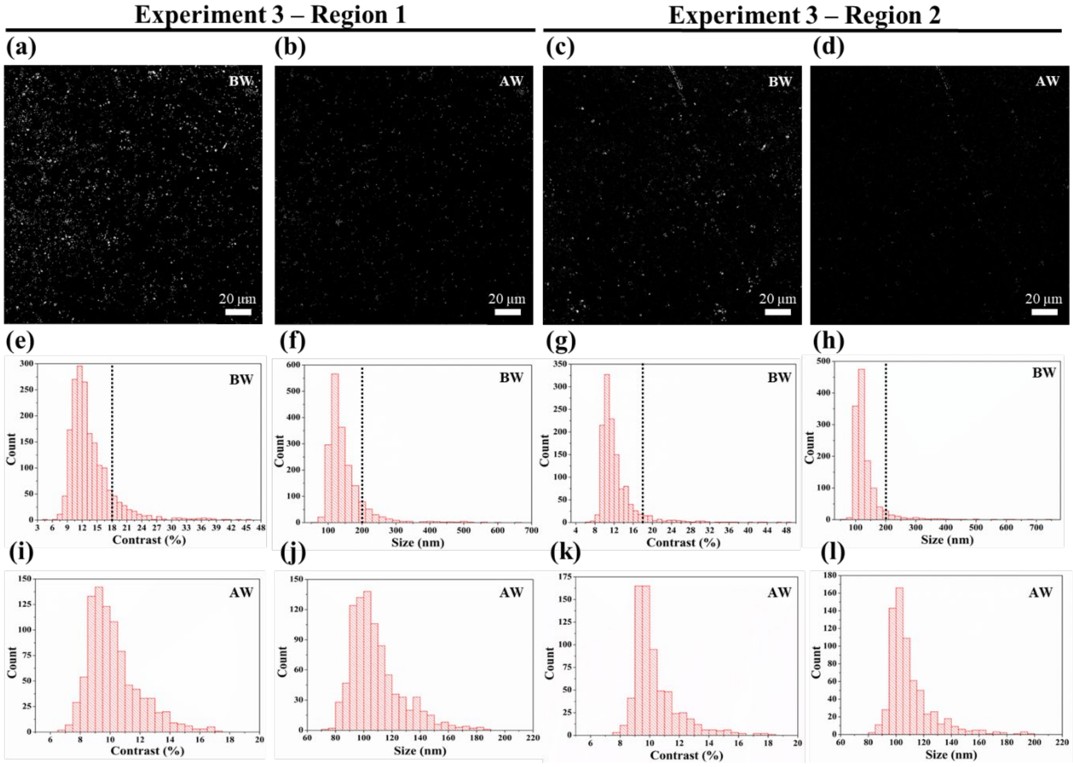


*Supplementary Figure 6: (a, c) PANORAMA images showing EV detection at 60 minutes (BW) in regions 1 and 2 for experiment 3. (b, d) PANORAMA images illustrating sEV detection (AW) in regions 1 and 2 for the same experiment. Histograms of EV contrast during BW are displayed in (e) region 1 and (g) region 2, alongside histograms of EV diameter in (f) region 1 and (h) region 2, with dashed lines indicating the cut-off threshold separating sEVs from larger EVs. Similarly, histograms of sEV contrast during AW are shown in (i) region 1 and (k) region 2, while histograms of sEV diameter are presented in (j) region 1 and (l) region 2, highlighting the size distribution of retained sEVs.*

### Supplementary Note 5:

Fabricating polydimethylsiloxane (PDMS) was used for producing our microfluidic device. The process involves casting PDMS onto a patterned SU-8 mold on a silicon wafer template. The following steps outline the fabrication of PDMS channel flow cells (Supplementary Fig. 7):

a) Clean a 4-inch silicon wafer with acetone, deionized water (DI), and isopropanol (IPA) to remove contaminants.

b) Spin-coat SU-8 photoresist on the wafer at 2000 rpm for 30 seconds. Perform a soft bake at 65°C for 5 minutes, followed by 95°C for 20 minutes.

c) Expose the SU-8 to UV light at 250 mJ/cm² for 10 seconds (lamp power 25 mW/cm²), followed by post-exposure bake at 65°C for 5 minutes, then 95°C for 10 minutes.

d) Develop the SU-8 pattern in developer for 10 minutes, rinse with DI water, dry with nitrogen gas, and perform a hard bake at 150°C for 10 minutes.

e) Mix PDMS (10:1 ratio of base to curing agent) and pour over the SU-8 mold. Cure at 70°C for 2 hours.

f) Peel the cured PDMS, treat with oxygen plasma at 50 W for 30 seconds to enhance hydrophilicity.

g) Transfer the PDMS channel flow cells onto arrayed gold nanodisk substrates (AGNIS) for further use.

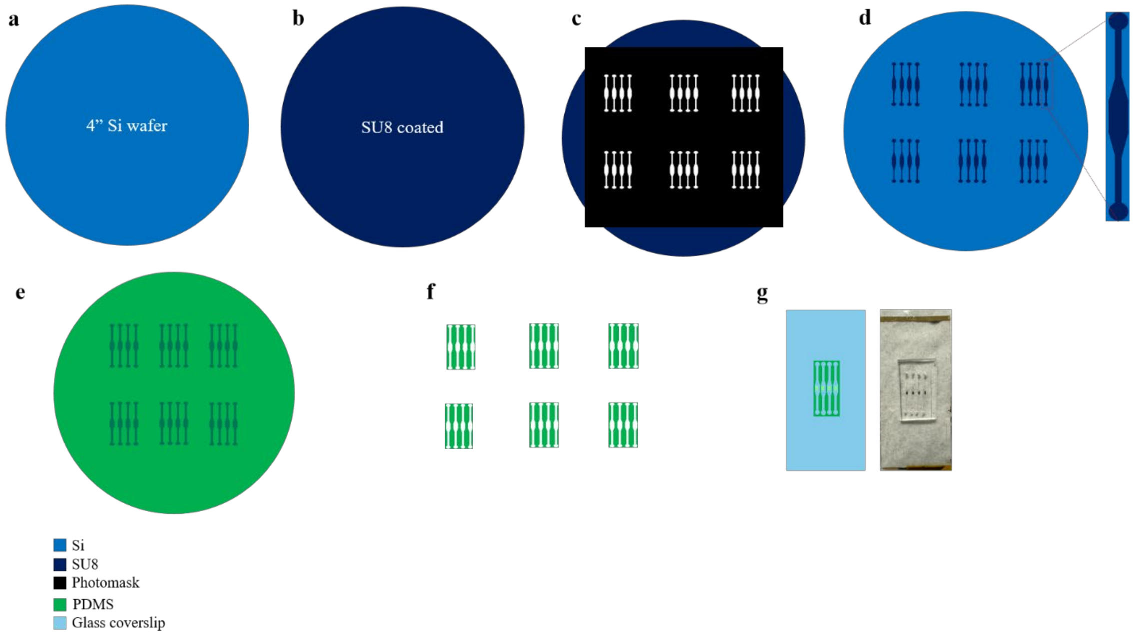


*Supplementary Figure 7: Fabrication steps of PDMS flow cell*
